# Auditory prediction hierarchy in the human hippocampus and amygdala

**DOI:** 10.1101/2022.11.16.516768

**Authors:** Athina Tzovara, Tommaso Fedele, Johannes Sarnthein, Debora Ledergerber, Jack J. Lin, Robert T. Knight

**Author notes:** **Corresponding author** Athina Tzovara University of Bern, Institute for Computer Science Neubrückstrasse 10, CH-3012 Bern.

## Abstract

Our brains can extract structure from the environment and form predictions given past sensory experience. Predictive circuits have been identified in wide-spread cortical regions. However, the contribution of subcortical areas, such as the hippocampus and amygdala in the formation of predictions remains under-explored. Here, we hypothesized that the hippocampus would be sensitive to predictability in sound sequences, while the amygdala would be sensitive to unexpected violations of auditory rules. We presented epileptic patients undergoing presurgical monitoring with standard and deviant sounds, in a predictable or unpredictable context. Onsets of auditory responses and unpredictable deviance effects were detected at earlier latencies in the temporal cortex compared to the amygdala and hippocampus. Deviance effects in 1-20 Hz local field potentials were detected in the lateral temporal cortex, irrespective of predictability. The amygdala showed stronger deviance responses in the unpredictable context. Additionally, low frequency deviance responses in the hippocampus (1-8 Hz) were observed in the predictable but not in the unpredictable context. Our results reveal a distributed cortical-subcortical network underlying the generation of auditory predictions, comprising temporal cortex and the hippocampus and amygdala, and suggest that the neural basis of sensory predictions and prediction error signals needs to be extended to subcortical regions.

## Introduction

The human brain has an astonishing capacity in detecting patterns from the environment in a quick and automatic way (Bar 2009). Detecting patterns in sensory stimuli allows making predictions about future events before they occur, based on current sensory input (Heilbron and Chait 2018). Every time that a pattern is violated, an internal model of the world is updated through prediction error signals (PE), which quantify the difference between expected and received outcome (Heilbron and Chait 2018). One experimental testbed for studying sensory predictions is through auditory deviance paradigms. These paradigms comprise of series of commonly repeated (standard) sounds, which are occasionally replaced by deviant tones (Garrido et al. 2009; Tivadar, Knight, and Tzovara 2021).

Because of the difficulty in assessing electrophysiological activity in subcortical regions in a non-invasive way, the search for a sensory predictive network in the auditory modality has mainly focused on a 2-node circuit, including mainly the temporal lobe and prefrontal areas (Garrido et al. 2009; Dürschmid et al. 2018; 2016; Rosburg et al. 2005; Canolty et al. 2006; Phillips et al. 2016a). The most prevalent view is that sensory areas compute a low-level predictive signal, comparing current sensory input to the immediate past and detect violations of auditory sequences (Dürschmid et al. 2016). This 2-node temporal-to-prefrontal circuit underlying sensory predictions has been well characterized by non-invasive (Garrido et al. 2008; Chennu et al. 2013) and invasive electrophysiology (Dürschmid et al. 2016; Rosburg et al. 2005; Phillips et al. 2016a; Edwards et al. 2005), establishing a cortical hierarchy of auditory predictions.

Both the hippocampus and amygdala are sensitive to auditory stimuli (Cusinato et al. 2022) and infrequent auditory events (Halgren et al. 1980; Knight 1996), but their specific role in the formation of auditory predictions remains unclear, as well as their integration with other cortical areas (E. Johnson et al. 2020). Previous studies based on functional Magnetic Resonance Imaging (fMRI) or Magnetoencephalography (MEG) have shown that the hippocampus is sensitive to violations of expected events (Kumaran and Maguire 2006a; Chen et al. 2013; Garrido et al. 2015), mainly through low frequency oscillations (Garrido et al. 2015; Recasens, Gross, and Uhlhaas 2018). Intriguingly, the hippocampus is also sensitive to unexpected visual events (Axmacher et al. 2010), and memory functions (E. L. Johnson and Knight 2015), possibly arbitrating between predictions and encoding of memories (Sherman and Turk-Browne 2020).

The amygdala is also sensitive to unexpected novel events (J. Blackford et al. 2010; Balderston, Schultz, and Helmstetter 2013), and to violations of expected auditory input (James et al. 2012). Interestingly, invasive electrophysiology recordings in macaques showed that single unit activity in the amygdala is sensitive to deviant auditory stimuli, with comparable latencies to those of prefrontal neurons (Camalier et al. 2019). Despite ample evidence for the involvement of the hippocampus and amygdala in detecting violations of environmental regularities, the specific function of each region in detecting deviant inputs and prediction remains underexplored.

Here, we aimed at shedding light on the role of the hippocampus and amygdala in detecting violations of auditory rules, and contrasting that to the well-established role of the temporal cortex (Dürschmid et al. 2016; Rosburg et al. 2005; Edwards et al. 2005). We hypothesized that in addition to a cortical hierarchy in processing auditory events, there also exists a sub-cortical one, including the hippocampus and amygdala. To dissociate effects of deviance and predictability, we used a paradigm comprising of standard and deviant sounds, presented in a temporally predictable or unpredictable way (Dürschmid et al. 2016). We recorded intracranial electroencephalography (iEEG) in patients with epilepsy to directly assess neural activity of the hippocampus and amygdala. We provide evidence for a distributed cortical-subcortical network underlying the generation of auditory predictions and highlight the role of the hippocampus and amygdala in detecting auditory rules and their violations.

## Materials and Methods

### Patients

We recorded intracranial EEG data in eight patients with pharmacoresistant epilepsy (mean age: 29 y, 3 women), undergoing pre-surgical monitoring. All patients had implanted depth electrodes, targeting the hippocampus and amygdala, among other regions (Supplemental Table 1 for an overview of electrodes across patients). Recordings took place at the University of California Irvine Medical Center, USA and at the University of Zurich (implantation), and the Swiss Epilepsy Center in Zurich (recordings), Switzerland. Patients gave written informed consent to participate in this study, approved by institutional ethics review boards of the University Hospital of Zurich (PB 2016–02055), UC Berkeley, and UC Irvine. All experiments were performed in accordance with the 6th Declaration of Helsinki.

### Paradigm

Patients were presented with series of standard (80%) and deviant (20%) sounds. Sounds were pure tones, lasting 100 ms. The standard sounds’ pitch was drawn from a gaussian distribution with μ=500 Hz σ2 = 125 Hz. The pitch for deviant sounds was at the tail of the distribution with a frequency of 2000 Hz. Deviant sounds were presented in a temporally predictable (after 4 standards) or unpredictable (after 3-8 standards) context (Figure 1). The sound-to-sound interval was 600 ms. Sounds were presented in two blocks of 500 trials, one for the predictable and one for the unpredictable context, lasting approximately 5’ each. Two additional blocks were recorded, in which the pitch of the standard sounds was drawn from a distribution with high variance, but were not analyzed, as they were out of the scope of the present study. The order of blocks was randomized for each patient. Patients were instructed to watch a silent video and ignore the sounds.

**Figure 1.**
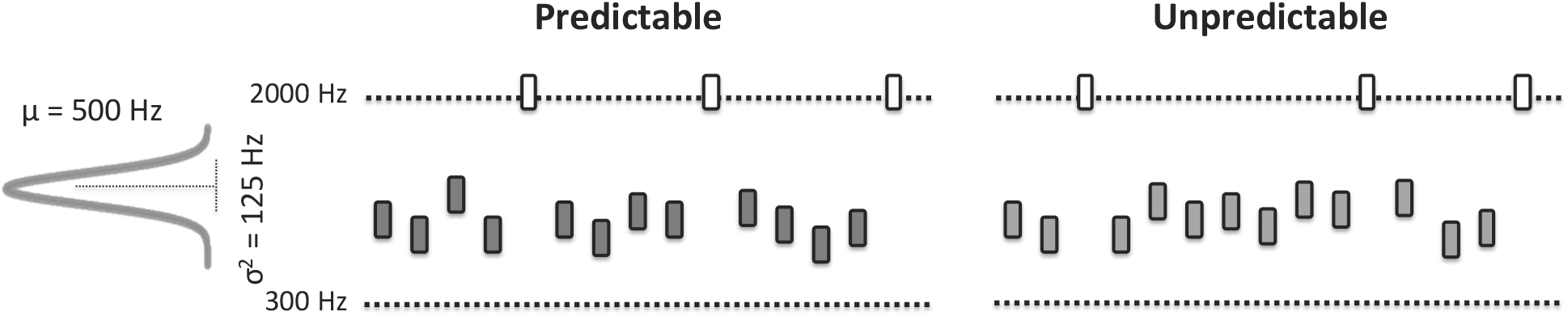
Task design. Patients were presented with series of standard (80%, gray) and deviant (20%, white) sounds. The standard sounds’ pitch was drawn from a gaussian distribution with μ=500 Hz σ^2^ = 125 Hz. Deviant sounds laid at the tail of the distribution with a frequency of 2000 Hz. Deviant sounds were presented in a temporally predictable (after 4 standards) or unpredictable (after 3-8 standards) context.

### Acquisition and pre-processing of electrophysiological data

Intracranial EEG was recorded over arrays of depth electrodes, typically consisting of eight stainless contacts each (AD-Tech, electrode diameter: 3 mm, inter-contact spacing: 10 mm). Contact location was identified by co-registering a post-operative computed tomography scan (CT) with a pre-operative high resolution anatomical magnetic resonance image (MRI). By considering the most consistent localizations of contacts across all patients, we analysed responses within 3 regions of interest, covering the hippocampus, amygdala and temporal cortex (TC) (Supplemental Table 1).

### Electrode localization

Electrodes were localized using merged post-operative computed tomography (CT) and pre-operative structural T1-weighted MRI scans. The CT scan was registered to the pre-operative MRIs, using a standard electrode localization procedure, implemented in Fieldtrip (Stolk et al. 2018). The electrode locations were visualized for each patient in native space, and their location was identified by a trained neurologist. To visualize the electrode locations across the group of patients, the aligned electrodes were warped onto a template brain in MNI space. For each patient, we retained for further analysis MTL electrodes which were localized in three different subregions (hippocampus / amygdala / temporal cortical areas).

### Data pre-processing

All data were visually inspected by a neurologist (RTK) to (a) exclude electrodes that were within the seizure onset zone, and (b) exclude periods of epileptic activity in the remaining electrodes. Continuous data were notch filtered, down-sampled to 500 Hz and re-referenced to a bipolar montage, according to their nearest neighbor on the same depth, to remove any source of noise from the common reference signal, following recommendations in the analysis of iEEG data (Lachaux et al. 2012; Mercier et al. 2022). All electrodes were band-pass filtered between 0.1 and 20 Hz prior to the extraction of local field event related potentials.

Peri-stimulus epochs were then extracted, spanning from -100 ms before the sounds’ onset to 500 ms post-stimulus onset. All epochs were then visually inspected to exclude any remaining artifacts. Data processing was performed using MNE python (Gramfort et al. 2013). For analyzing local field potentials, epochs were baseline corrected with the mean of a pre-stimulus baseline (−100 to 0 ms), which was randomly chosen from the pool of baseline time points of all trials, based on 500 iterations, similar to previous work with iEEG data (Kam et al. 2019).

### Responsive electrodes

As the electrode placement was driven by clinical criteria, we focused all analyses on electrodes that responded to the auditory stimulation. To not bias our search for deviance and predictability effects, we pulled all cleaned epochs together and sought electrodes that responded to all sounds irrespective of the sounds’ identity.

For each electrode, we contrasted the time-point by-time post-stimulus local field potentials with the mean of the pre-stimulus activity (−100 to 0 ms), using t-tests. The resulting t-values were corrected for multiple comparisons based on the false discovery rate (p<0.05). Onsets of responsiveness were defined at the single electrode level, by considering the first time point that showed a significant response, while peaks by considering the location of the maximum absolute value among all significant t-values. Onset and peak latencies were contrasted at the group level, pulling all electrodes together across regions using linear mixed effect models, with a random intercept of patient to account for across-patient differences.

### Deviance effects

Deviance effects were parametrized by F-values, which were computed for each electrode, and context based on a 1-way anova, with a factor of deviance, as in a previous studies using similar paradigms (Kam et al. 2021; Dürschmid et al. 2016). Onsets of deviance effects for each context were defined as the first time-point where significance was reached, assessed by comparing the true values to the distribution of effects obtained via 1000 random permutations in the labels of standard and deviant epochs (Dürschmid et al. 2016).

Group-level effects of predictability were identified by further contrasting the time-courses of F-values quantifying deviance effects for the predicable and unpredictable contexts, as in previous studies using a similar paradigm (Dürschmid et al. 2016). This approach is equivalent to a 2 by 2 Anova, and was preferred over a ‘ classical’ implementation of a 2-factorial test, because of the bipolar reference in the data: as electrodes were re-referenced to their neighbors, a positive peak in one of them could deflect as a negative peak in the next, and close to zero on average. By parametrizing LFPs with F-values, the sign of LFP measures becomes irrelevant, and only information about the magnitude of effects is retained, making it possible to perform group-level analyses.

### Time-frequency analysis

For each responsive channel we decomposed the time course of LFPs in time-frequency representations, using Morlet wavelets with multi-taper windows (function tfr_array_multitaper from MNE), applied with 0.5 Hz steps. The resulting single-trial power was normalized by the log-ratio of the pre-stimulus baseline. Singletrial power was then averaged within experimental conditions.

### Statistical contrasts

Statistical tests grouping data from multiple patients, for example testing for response onsets, were based on linear mixed effects models, with a random intercept for accounting for different patients, as it is common practice in the field (Cusinato et al. 2022; E. L. Johnson et al. 2018). Bonferroni correction was used for correcting for multiple comparisons. For statistical tests performed in time frequency analyses of iEEG signals correction for multiple comparisons was achieved via cluster-based permutation tests (p<0.05, 1000 permutations).

## Results

Patients were presented with series of standard and deviant sounds (80% and 20% of the time, respectively). Deviant sounds were presented in a temporally predictable (i.e. always after five standard sounds), or unpredictable (after 3-8 standard sounds) context (Figure 1). Patients were instructed to focus their attention to a silent movie and ignore the presented sounds.

### Responsive electrodes

We first assessed electrodes that were responsive to all sounds, irrespective of context and deviance manipulations, by testing for significant changes in 1-20 Hz local field event related potentials (LF-ERPs) with respect to a 100 ms baseline period. Patients had consistent electrode implantations in the lateral and medial temporal lobe, covering the lateral temporal cortex and the hippocampus and amygdala (Supplemental Table 1 for electrode coverage per patient and region). All three regions contained responsive electrodes to sounds across patients (Supplemental Table 1). Electrodes in the temporal cortex showed a significantly earlier response onset than electrodes in the amygdala (Figure 2a, F(1,43) = 8.08, p < 0.05, linear mixed effects models accounting for different patients here and in the following), and an earlier response than electrodes in the hippocampus (F(1,45) = 4.06, p < 0.05). The mean response onset across patients and electrodes was at 79 ms for the temporal cortex, 134 ms for the hippocampus and 154 ms for the amygdala (Figure 2a).

**Figure 2.**
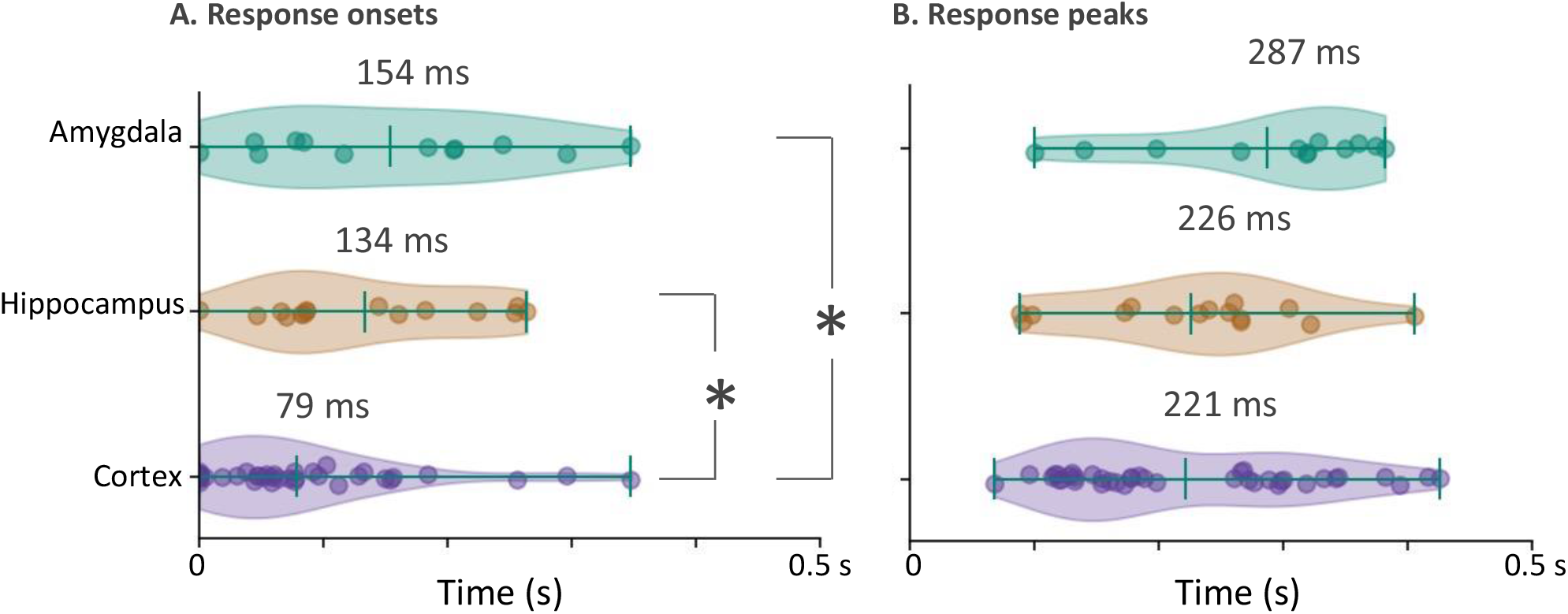
Onsets and peaks for responsive amygdalar, hippocampal and cortical electrodes in LF-ERPs (1-20 Hz). Responsiveness was assessed by merging responses to all sound categories. Responsive electrodes in the cortex showed a significantly earlier onset compared to hippocampal and amygdalar ones (F(1,45) = 4.06, p < 0.05 for cortex vs. hippocampus and F(1,43) = 8.08, p < 0.05 for cortex vs. amygdala).

There was no significant difference between the onsets of the hippocampus and amygdala (F(1,18) = 0.12, p = 0.74). Response peaks showed a similar tendency as onsets, with peak responses occurring at 221 ms on average for temporal electrodes, and at 226 ms for hippocampal and 287 ms for amygdalar ones (Figure 2b).

### Deviance effects

Focusing on responsive electrodes, we then contrasted local field event related potentials (LF-ERPs) in response to standard vs. deviant sounds. As expected from previous studies (Dürschmid et al. 2016), these showed a strong deviance response in temporal areas (Figure 3 for exemplar LF-ERPs), and deviance responses for both the predictable and unpredictable contexts (Figure 3c/d for exemplar responses). The deviance effects in amygdala and hippocampus are shown in Figure 4. To quantify deviance effects at the group level, we parametrized LF-ERPs in response to standard vs. deviant sounds for each context, by F-values, which quantify the strength of deviance effects (Figure 5a for group level deviancy effects). Group level results in Figure 5 are visualized via F-values similar to previous studies (Dürschmid et al. 2016), and not as LF-ERPs. Because a bipolar reference was used in the analysis, neighbor electrodes can have responses of opposing sign, and therefore averaging all LF-ERPs is not meaningful. F-values overcome this issue, as they quantify the strength of deviance effects across trials, irrespective of the sign of LF-ERPs responses. At the group level, the temporal cortex showed the strongest deviance effects for both predictable and unpredictable contexts (Figure 5a, purple plots). However, there was no significant difference in deviance responses between the two contexts.

**Figure 3.**
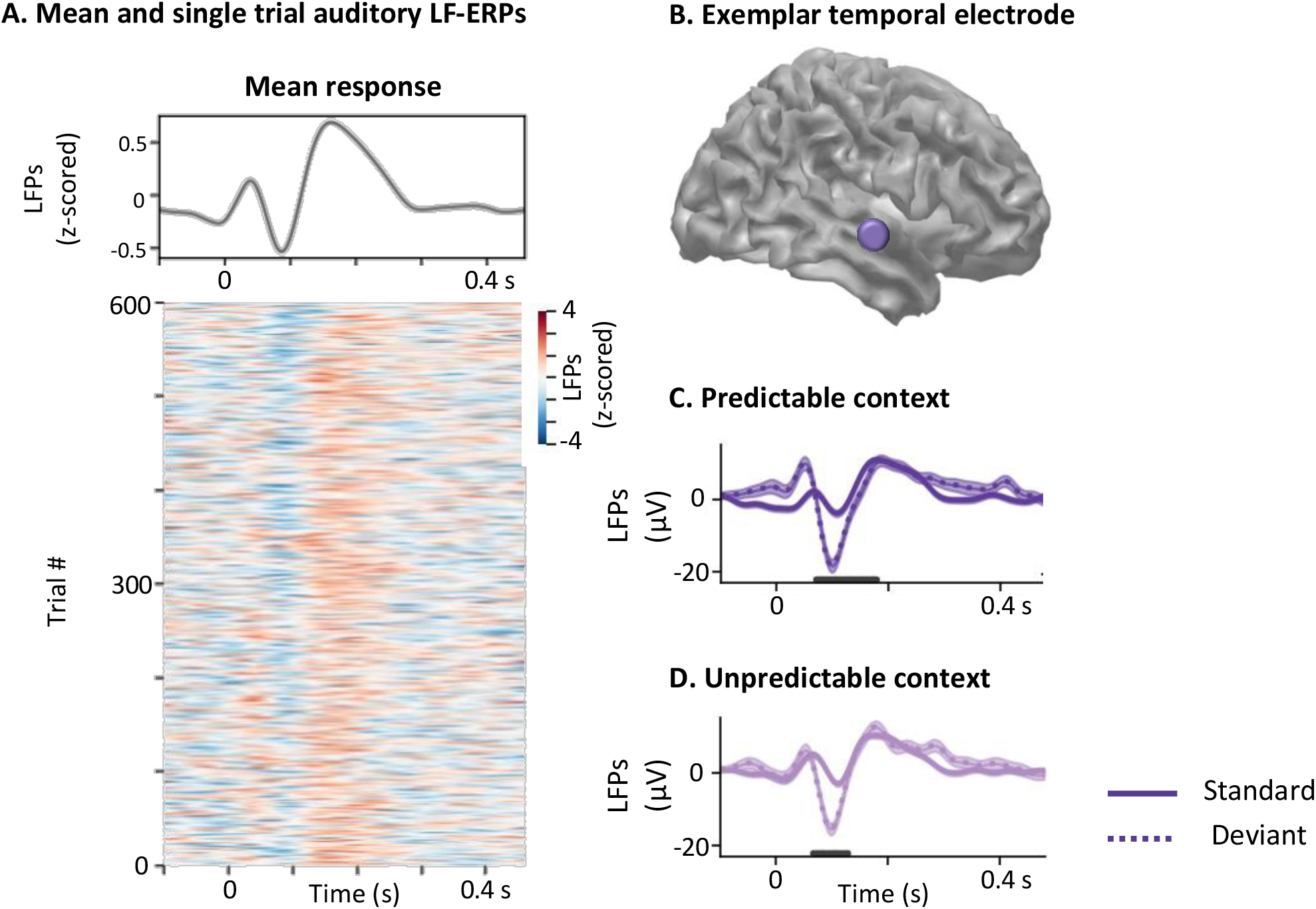
Exemplar LF-ERPs in the temporal cortex. A. Mean and single trial LF-ERPs for an exemplar electrode in the temporal cortex. B. Location of this electrode with MNI coordinates: -56.79, -16.11, -4.90. C/D. Responses to standard (full) vs. deviant (dotted lines) sounds for the predictable (C) and unpredictable (D) contexts. Horizontal lines highlight periods of significant difference between standard and deviant responses.

**Figure 4.**
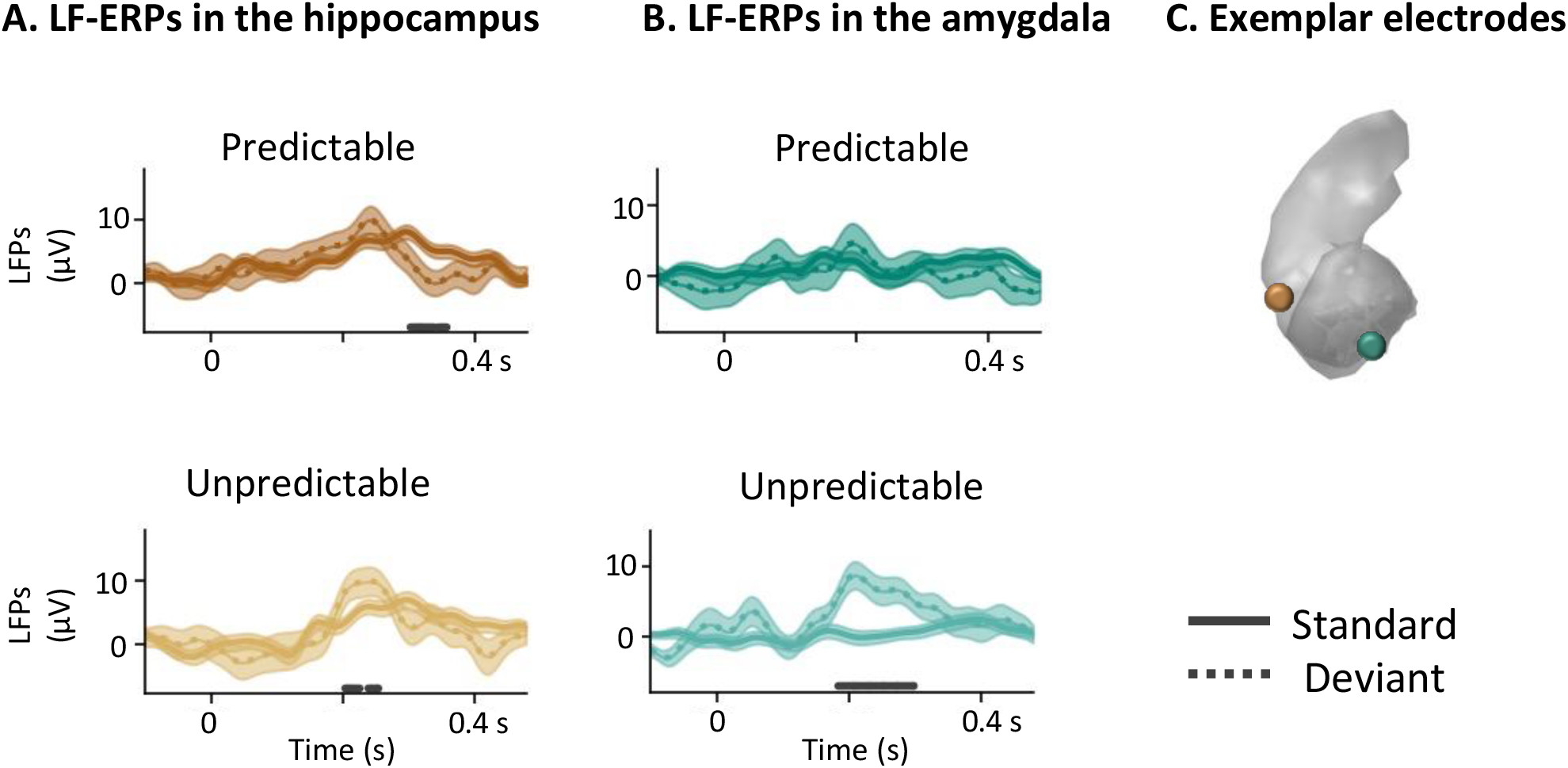
LF-ERPs in the hippocampus (A) and amygdala (B). Top rows represent 1-20 Hz LF-ERPs in response to predictable and bottom rows to unpredictable standard and deviant stimuli. Full lines show responses to standard and dotted to deviant sounds. Horizontal lines highlight periods of significant difference. Although the hippocampus showed a deviance response both for predictable and unpredictable contexts, the amygdala showed a deviance response for the unpredictable context only. Horizontal lines highlight periods of significant deviance effects, assessed through permutation statistics. C. Location of this contact, MNI coordinates: [37.51, -12.92, -19.24] for the hippocampus and [-21.99, -1.19, -24.26] for the amygdala.

**Figure 5.**
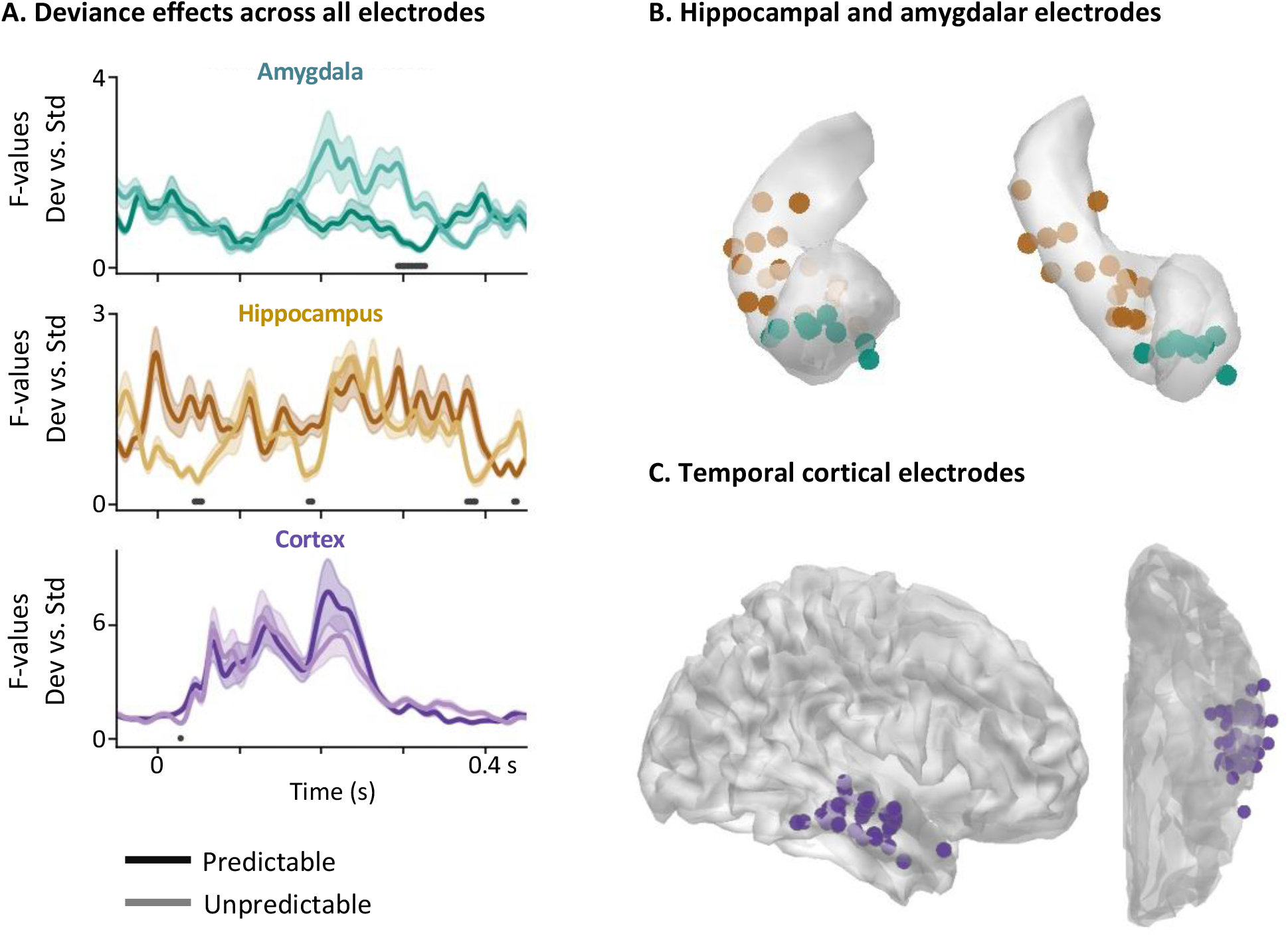
Group level deviance effects. A. Time-course of deviance effects for predictable (dark lines) and unpredictable (light lines) contexts across all contacts and patients. Deviance effects shown in panel (A) are quantified via F-values, computed by contrasting LF-ERPs in response to standard vs. deviant sounds at the single-trial level, similar to previous studies (Dürschmid et al. 2016). The amygdala (top row) showed a stronger deviance response for the unpredictable context compared to predictable one, while the hippocampus for the predictable context. By contrast, deviance responses in the cortex did not differ between the two contexts. Horizontal black lines highlight periods of significant difference in F-values for predictable vs. unpredictable contexts. B/C. Overview of all electrodes for hippocampus, amygdala (B) and cortex (C), projected on MNI templates.

The hippocampus by contrast, showed deviance responses in the LF-ERP range for both contexts (Figure 4a for exemplar LF-ERPs in the hippocampus). At the group level, we observed a significantly stronger deviance effect for the predictable context compared to the unpredictable one (Figure 5a, for group results quantified via F-values, black marks on x-axis denote significant differences between predictable and unpredictable contexts). In the amygdala by contrast, the strongest deviance effects were observed for the unpredictable context, and at late latencies, around 300 ms post-stimulus onset (Figure 5a for group F-values, Figure 4b for exemplar LF-ERPs).

### Latencies of deviancy effects

After evaluating whether deviancy effects within each region of interest are modulated by predictability, we additionally characterized their latency for each of the two contexts separately, at the level of single electrodes. These latencies refer to the onset of deviance responses, i.e. differences between standard and deviant sounds in the two contexts, and not to the latency of an auditory -sensory-response (which is shown in Figure 2). In the unpredictable context, the temporal cortex showed the earliest deviance effects, with a mean onset across patients and contacts of 100 ms (Figure 6, light colors). Deviance effects in the hippocampus had mean onset at 210 ms (Figure 6, light colors), significantly later than the temporal cortex (F(1,39)=12.83, p_corr_ <0.01). In the amygdala, unpredictable deviance had a mean onset at 195 ms, which was not significantly later than the temporal cortex when correcting for multiple comparisons (F(1,36)=7.61, p_corr_ =0.055).

**Figure 6.**
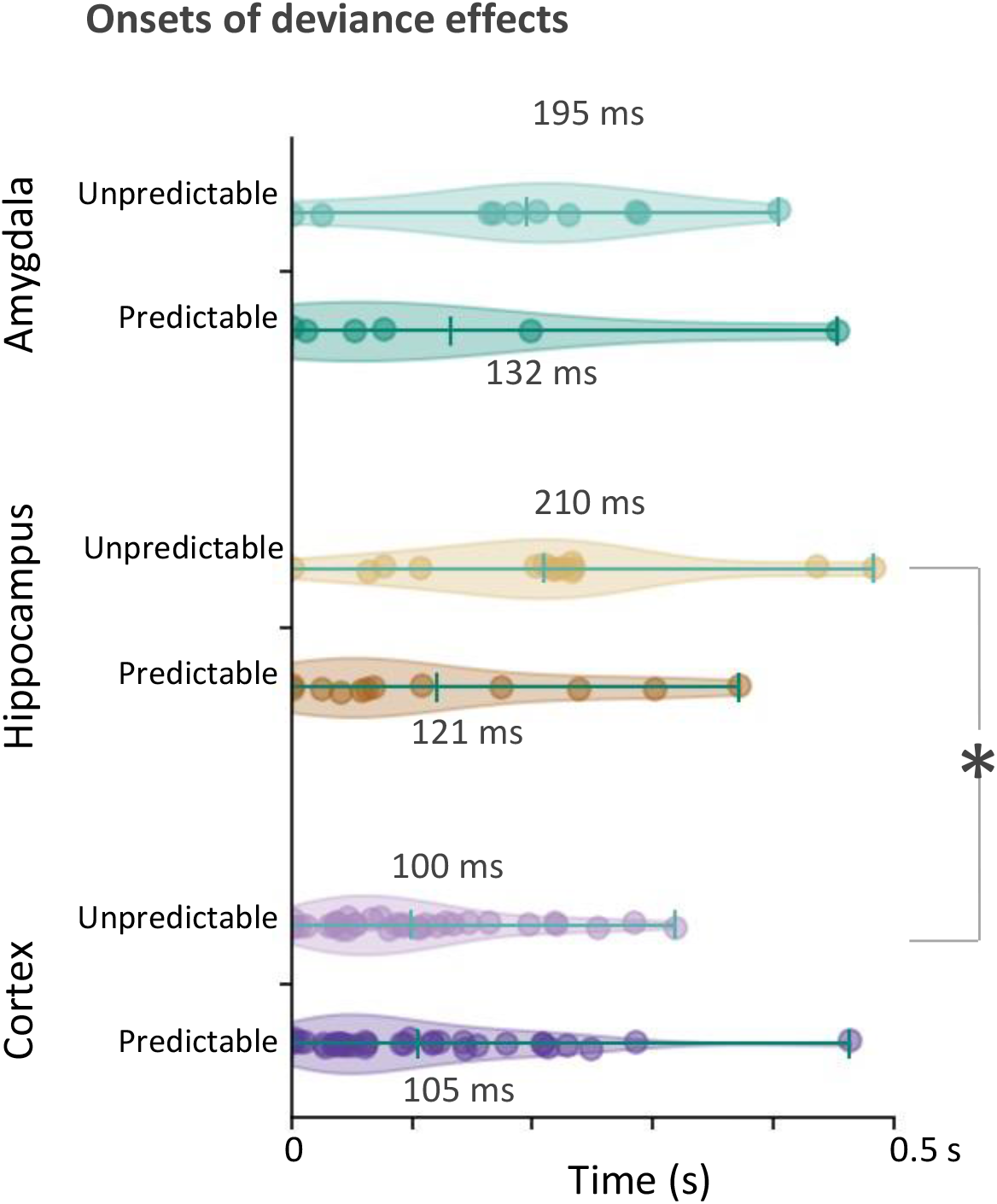
Onsets of deviance effects in LF-ERPs for individual contacts split by context. For the unpredictable context, the onsets of deviance effects occurred at earlier latencies for temporal compared to hippocampal electrodes (F(1,39)=12.83, pcorr<0.01). No difference was observed for the predictable context.

For the predictable context, the temporal cortex had shorter latencies (105 ms post-stimulus onset) than the hippocampus (121 ms) and amygdala (132 ms), but these were not significantly different (Figure 6, dark plots). Deviancy onset effects in the hippocampus were shorter for the predictable (121 ms) compared to the unpredictable (210 ms) contexts, but this difference was not statistically significant when correcting for multiple comparisons (F(1,18)=4.17, p_corr_ =0.17, Figure 6).

### Frequency contents of deviance responses

We next evaluated the frequency content of local field potential responses to the auditory stimuli, by computing time-frequency analyses for the three regions of interest at the group level (Figure 7). These revealed a significant effect of deviance only in the hippocampus (Figure 7). Low frequency power, in the range of 1-8 Hz was significantly stronger in response to deviant compared to standard sounds in the hippocampus, for the predictable context only, for a sustained period starting around 100 ms post-stimulus onset (Figure 7, gray outline highlighting a significant cluster, with a cluster-level p_corr_ <0.05). Higher low-frequency power for deviant sounds compared to standards was observed for all patients for the predictable, but not unpredictable context (Supplemental Figure 1). In the cortex and amygdala, there was a tendency for higher power in response to deviant sounds, but this was not significant after correcting for multiple comparisons.

**Figure 7.**
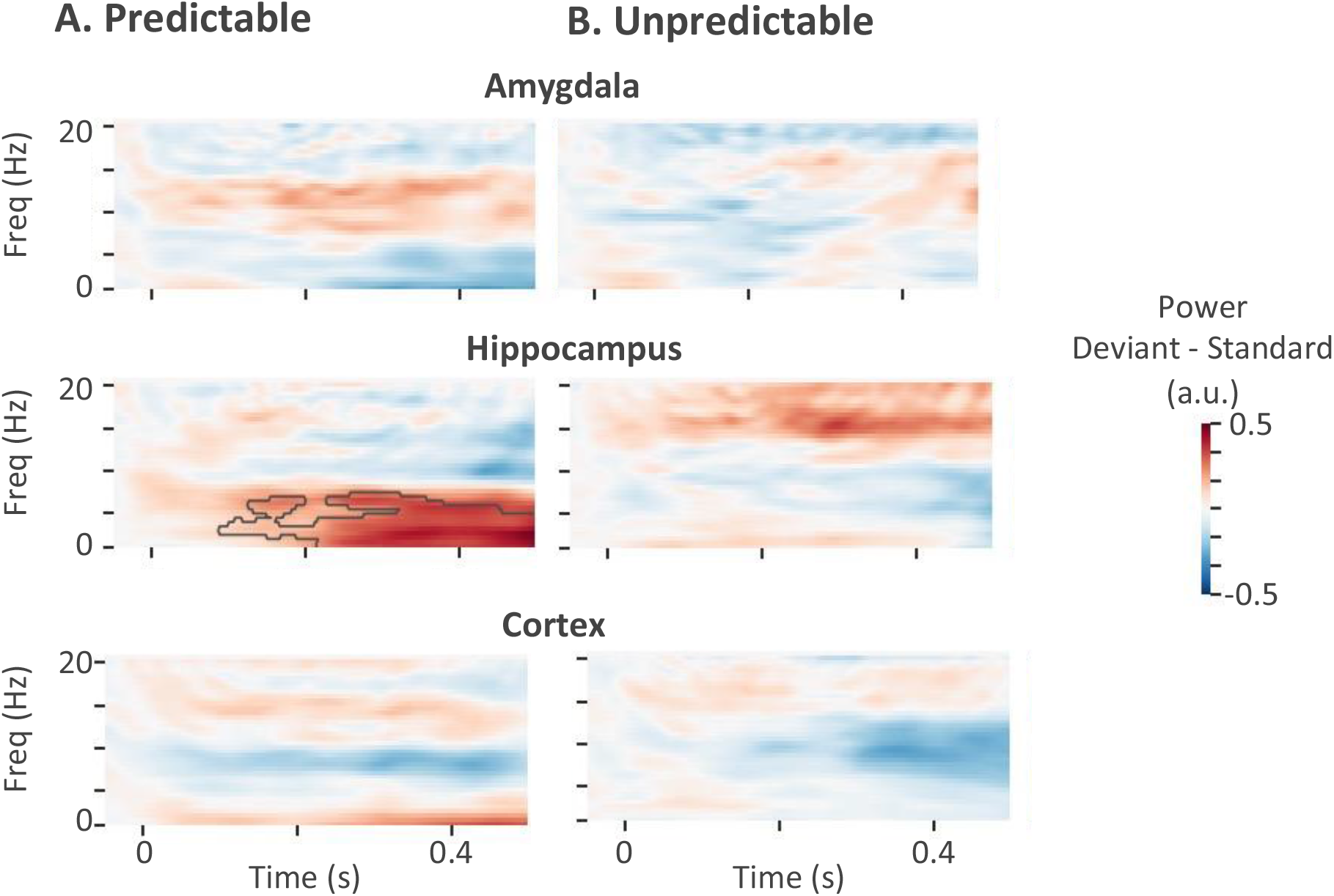
Deviant - standard power across patients and regions. Each plot illustrates the difference in average power in response to deviant – standard sounds, for the predictable (A) and unpredictable (B) contexts. A significant increase in low frequency power was observed for the predictable context only for the hippocampus (cluster-level p_corr_ <0.05).

### Low frequency activity in the hippocampus supports auditory predictions

Thus far we considered all standard sounds as one condition, irrespective of their temporal order of appearance. Next, we evaluated the link between low frequency activity in the hippocampus and auditory predictions. We focused on the frequency range that showed a deviance effect for the predictable context in the hippocampus (Figure 7) and we used it as mask to compute the average hippocampal power as a sequence unfolds. For this analysis, we split standard sounds into sub-groups, according to their order of presentation (Figure 8, S1-S4).

**Figure 8.**
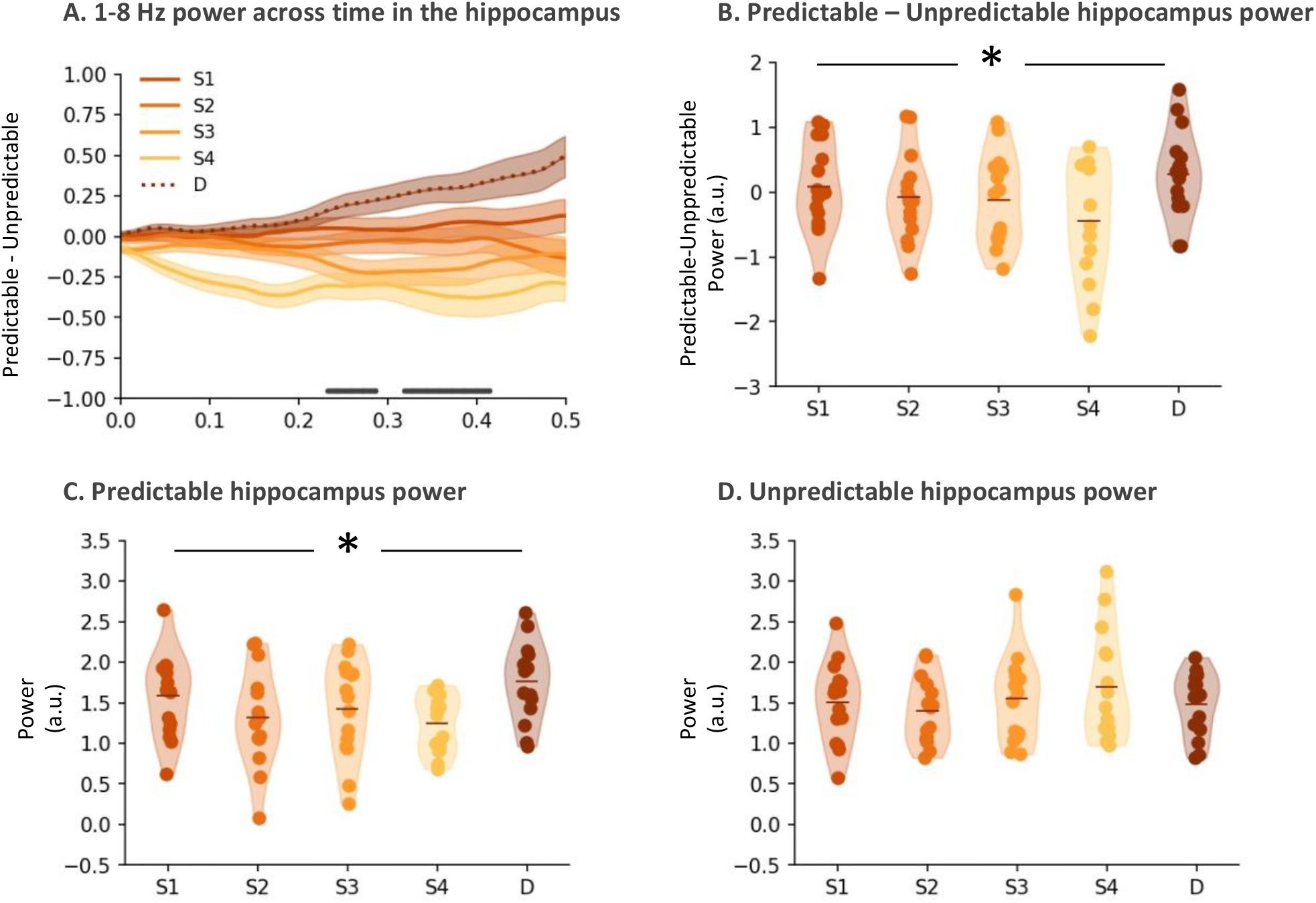
Low frequency power in the hippocampus as the auditory sequence unfolds. A. Full lines illustrate mean 1-8 Hz power in response to standard sounds, by order of presentation (S1-S4), subtracting power in response to predictable from unpredictable sounds. Dashed lines show 1-8 Hz power for deviant predictable minus unpredictable sounds. Horizontal lines highlight periods where the ordering was significant (p < 0.05). (B)Mean power in the hippocampus as the auditory sequence unfolds, for the predictable minus unpredictable context. The mean power was computed for each contact via the mask of significant deviance effects in hippocampus (Figure 7), and showed a significant effect of sequence (F(1,14) = 7.87, p_corr_ < 0.05). C/D. Mean power masked by deviance effects for the predictable (C) and unpredictable contexts (D) separately. Only the predictable context showed a significant effect of sequence (F(1,14) = 8.29, p_corr_ < 0.05).

We observed a significant effect of sequence in the difference of low frequency hippocampal power between predictable and unpredictable contexts (F(1,14) = 7.87, p_corr_ < 0.05, Figure 8b). This was driven by a gradual decrease of low frequency power as the predictable sequence unfolded, followed by a strong increase in response to the predictable deviant sound (Figure 8c, F(1,14) = 8.29, p_corr_ < 0.05). Post-hoc analysis revealed a significant difference between low frequency power in response to the deviant sound and the second or fourth standard sound in the sequence (p < 0.01, Figure 8c, S2 vs. D and S4 vs. D). Although there was a tendency for lower power between the third standard sound and the deviant, this was not significant (p = 0.07, Figure 8c, S3 vs. D). Control analysis for the unpredictable context showed no effect of sequence Figure 8d, F(1,14) = 1.27, p = 0.26).

Last, we also assessed changes in hippocampal power over time as the sequence unfolds. To this aim, to keep one consistent frequency band over time, we computed power in at 1-8 Hz (Figure 8a). 1-8 Hz power in the hippocampus was significantly modulated by the order of sound presentation between 232 and 290 ms, and between 318 and 414 ms post sound onset (Figure 8a, horizontal lines, F(1,14) = 7.7.00, p<0.01 at 260 ms). As a control analysis, there was no power modulation by auditory sequence neither in the amygdala (F < 3.01, p> 0.09), nor the temporal cortex (F < 5.2, p > 0.06).

## Discussion

Intracranial EEG recordings in humans provided direct evidence of subcortical contributions in the formation of auditory predictions. We found that the hippocampus and amygdala are sensitive to deviant sounds with distinct roles. Deviance effects in LF-ERPs in the hippocampus were stronger for the predictable compared to the unpredictable context, with an opposite pattern in in the amygdala showing enhanced responses for unpredictable deviance. By contrast, deviance effects in the temporal cortex were not modulated by predictability, in accordance to previous reports using a similar paradigm in a different group of patients (Dürschmid et al. 2016). Taken together, our findings suggest the existence of a distributed network in the medial temporal lobe underlying sensory predictions: while the temporal cortex computes ‘ low’ level predictions, comparing each sensory input to the immediate past, the hippocampus maintains a longer memory trace of auditory patterns, spanning over sequences of sounds, and is particularly active when a violation of the sequence can be predicted.

### Subcortical network underlying auditory predictions

Our findings expand the network of deviance detection beyond a 2-node cortical network, which has been excessively studied using mainly non-invasive imaging (Chennu et al. 2013; Garrido et al. 2009). Invasive EEG recordings have been used to confirm this 2-node network (Dürschmid et al. 2016; Phillips et al. 2016b), and have also demonstrated additional regions sensitive to deviance, including the insula (Blenkmann et al. 2019), the nucleus accumbens (Durschmid et al. 2016), the hippocampus and the amygdala (Halgren et al. 1980; Camalier et al. 2019). Here, we focused on the hippocampus and amygdala and showed that the latencies of processing deviant sounds in the amygdala follow that of the hippocampus and temporal cortex, suggesting a cortical-subcortical hierarchy in detecting auditory deviance. The fact that the amygdala responds to deviance at later latencies than the temporal cortex is in accordance with a recent monkey study using single unit activity (Camalier et al. 2019). Moreover, the hierarchy in auditory response latency observed in this study irrespective of deviance effects, fits the latencies of auditory responses reported in a recent study using a different auditory paradigm and patients, where the temporal cortex showed earlier responses compared to both the hippocampus and amygdala (Cusinato et al. 2022).

Importantly, our findings suggest that the amygdala is mainly sensitive to unexpected deviance, as indicated by a higher number of contacts showing deviance effects for an unpredictable compared to predictable context, and a stronger overall deviance effect (Figure 5). These findings reinforce the role of the amygdala as a novelty detector (J. U. Blackford et al. 2010), or as being sensitive to events of high saliency (Fedele et al. 2020) which could be particularly relevant for cases of unpredictable deviance, as an unexpected change in the environmental statistics might signal danger (Balderston, Schultz, and Helmstetter 2013).

The sensitivity of the hippocampus to violations of sequences has been mainly examined in the context of ‘ oddball’ paradigms, in the auditory (Ioannides et al. 1995) or somatosensory (Hamada et al. 2004) modalities, where participants are asked to actively detect rare target stimuli in a stream of regularly repeated events. Our finding that the hippocampus is mainly sensitive to predictable deviant sounds fits with findings of previous studies which have shown hippocampal sensitivity to violations of predicted events. In the visual modality, the hippocampus has been found to be responsive to violations of an established sequence of events, rather than completely novel events (Kumaran and Maguire 2006b; Chen et al. 2013). Garrido and colleagues (Garrido et al. 2015) used a visual sequence of 4 objects, presented in a fixed, mismatch and unpredictable order in a MEG study. Using source reconstruction techniques, authors reported a higher theta power in the mismatch compared to the fixed or unpredictable conditions. Similar findings have also been reported in the auditory modality (Recasens, Gross, and Uhlhaas 2018), with additional evidence that hippocampal-to-cortical connectivity underlies the encoding of predictable sequences. Additionally, the hippocampus, together with the temporal and prefrontal cortex, were found to underlie the detection of predictable auditory sequences (Barascud et al. 2016). These results are in accord with our findings, as we show a significant increase in low frequency hippocampal power in response to predictable but not unpredictable violations of expected auditory events.

Overall, our findings suggest that low frequency hippocampal activity contributes to the detection of auditory sequences and formation of auditory predictions. Low frequency activity in the hippocampus has been previously shown to mediate memory (E. L. Johnson et al. 2018; Boran et al. 2019; Dimakopoulos et al. 2022) and predictions of future events (Sherman and Turk-Browne 2020). Our results expand the functions of low frequency hippocampal activity towards a role in maintaining an active model of environmental regularities.We found that the difference in hippocampal low frequency power between standard and deviant sounds increases the closer a sequence gets to a deviant sound, but only when the occurrence of a deviant sound can be predicted (Figure 8). One interpretation for these findings is that the hippocampus plays an active role in updating an environmental model, keeping track of an ongoing sequence as it rapidly evolves across a sequence of auditory events. Similar findings have been reported for the prefrontal cortex, which has also been shown to be sensitive to ongoing auditory sequences (Dürschmid et al. 2018). Whether this tracking of auditory patterns is an inherent property of the hippocampus, or driven by external input, such as the prefrontal cortex, remains to be investigated.

### Limitations and future directions

One main limitation of our study is the sparse electrode coverage: because of our primary goal to investigate hippocampal and amygdalar contributions in deviance detection and auditory predictions we focused on patients that had good coverage in the medial temporal lobe, with at least one hemisphere being seizure-free. As a consequence, in our patient cohort there were no patients with frontal depth electrodes or grids, which would have allowed investigation of hippocampal-amygdalo-prefrontal interactions. Future studies can profit from recent advances of high-precision MEG to reconstruct subcortical activity (Tzovara et al. 2019) and study how the medial temporal lobe network interacts with prefrontal regions. Another limitation of our study is that because of the tight timing in our experimental setup we were not able to assess metrics of functional connectivity among our three target regions, which typically rely on oscillatory coupling, that evolves over longer temporal intervals (Dimakopoulos et al. 2022). Our findings on the timing of onsets and peaks auditory responses and deviance effects provide a first indication on the information flow in the amygdalo-hippocampal-temporal network, that can be confirmed using longer sound intervals.

## Conclusions

We provide evidence for the existence of a subcortical hierarchy underlying auditory predictions. Our findings complement existing studies that have focused on the cortex and suggest that the search for sensory predictions and prediction error signals needs extension to subcortical regions. Importantly, our findings suggest the existence of a distributed network underlying the generation of auditory predictions, comprising cortical sensory areas, which compute a ‘ low’ -level prediction, the amygdala, which is sensitive to unexpected violations of streams of sensory information, and the hippocampus, which computes auditory predictions through low frequency activity.

---

**Supplemental Table 1.** Description of patients and electrode locations. For each patient we display the total number of electrodes that were located in each of our regions of interest (Column Total) and the number of electrodes that were responsive to auditory stimuli (Column Responsive). Only electrodes that were located in seizure free regions are included.

| Patient | Age | Sex | Amygdalar electrodes |  | Hippocampal electrodes |  | Cortical electrodes |  |
| --- | --- | --- | --- | --- | --- | --- | --- | --- |
|  |  |  | Total | Responsive | Total | Responsive | Total | Responsive |
| 1 | 24 | F | 5 | 0 | 5 | 3 | 4 | 2 |
| 2 | 28 | M | 6 | 2 | 3 | 2 | 5 | 4 |
| 3 | 20 | F | 3 | 2 | 0 | 0 | 4 | 2 |
| 4 | 28 | M | 4 | 1 | 7 | 3 | 13 | 8 |
| 5 | 57 | M | 3 | 1 | 4 | 2 | 4 | 3 |
| 6 | 29 | M | 3 | 0 | 5 | 1 | 0 | 0 |
| 7 | 26 | M | 4 | 3 | 11 | 3 | 14 | 14 |
| 8 | 21 | F | 5 | 3 | 3 | 1 | 7 | 6 |
| Total | - | - | 33 | 12 | 38 | 15 | 51 | 39 |

**Supplemental Figure 1.**
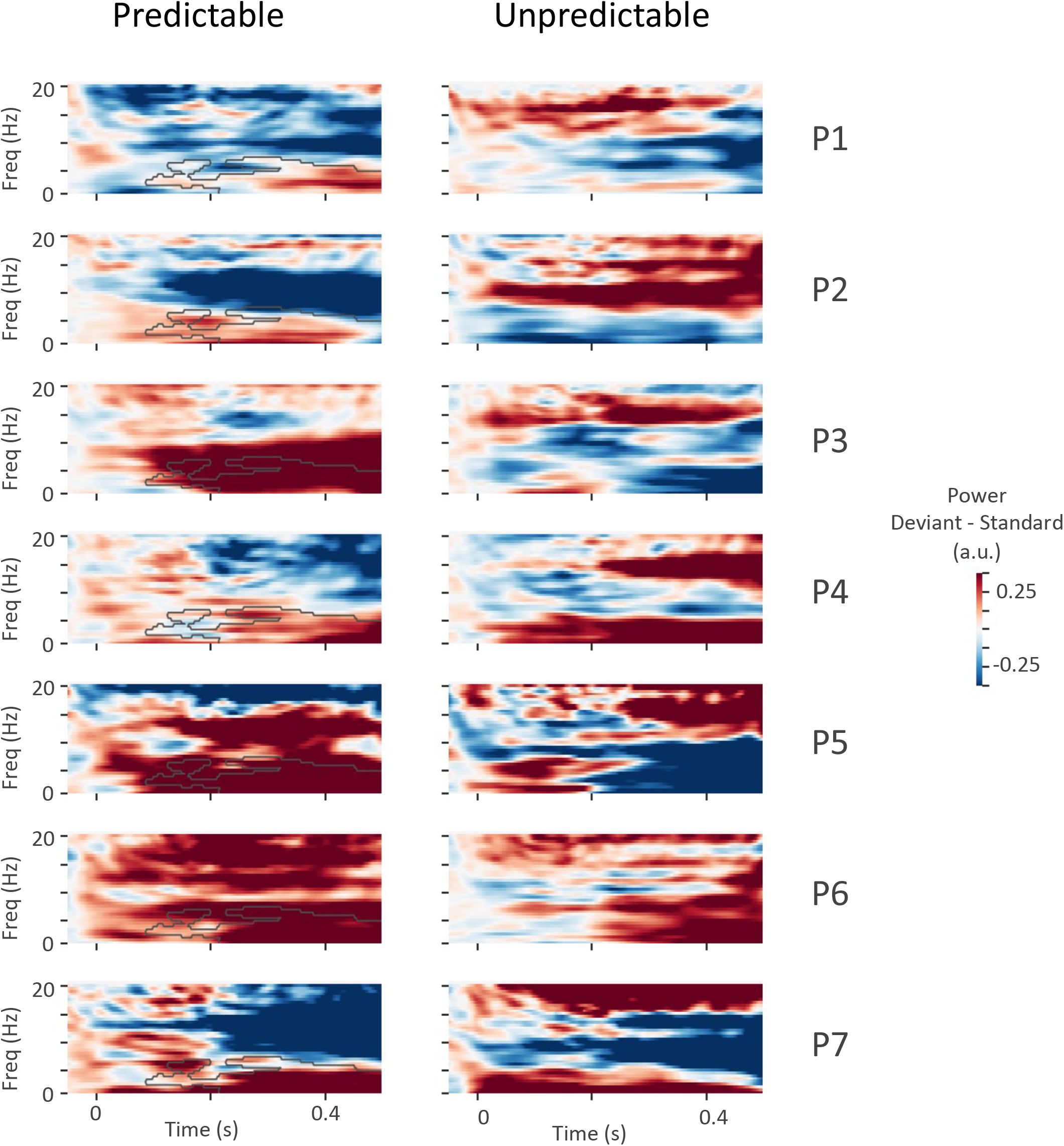
Single patient deviant - standard power in the hippocampus for the predictable (a) and unpredictable (b) contexts. For the predictable context, all patients had higher low frequency power in response to deviant vs. standard sounds. The plotted contour for predictable sounds highlights the boundaries of significant increase in low frequency power that was obtained at group level (Figure 7 of the main text).

**Supplemental Figure 2.**
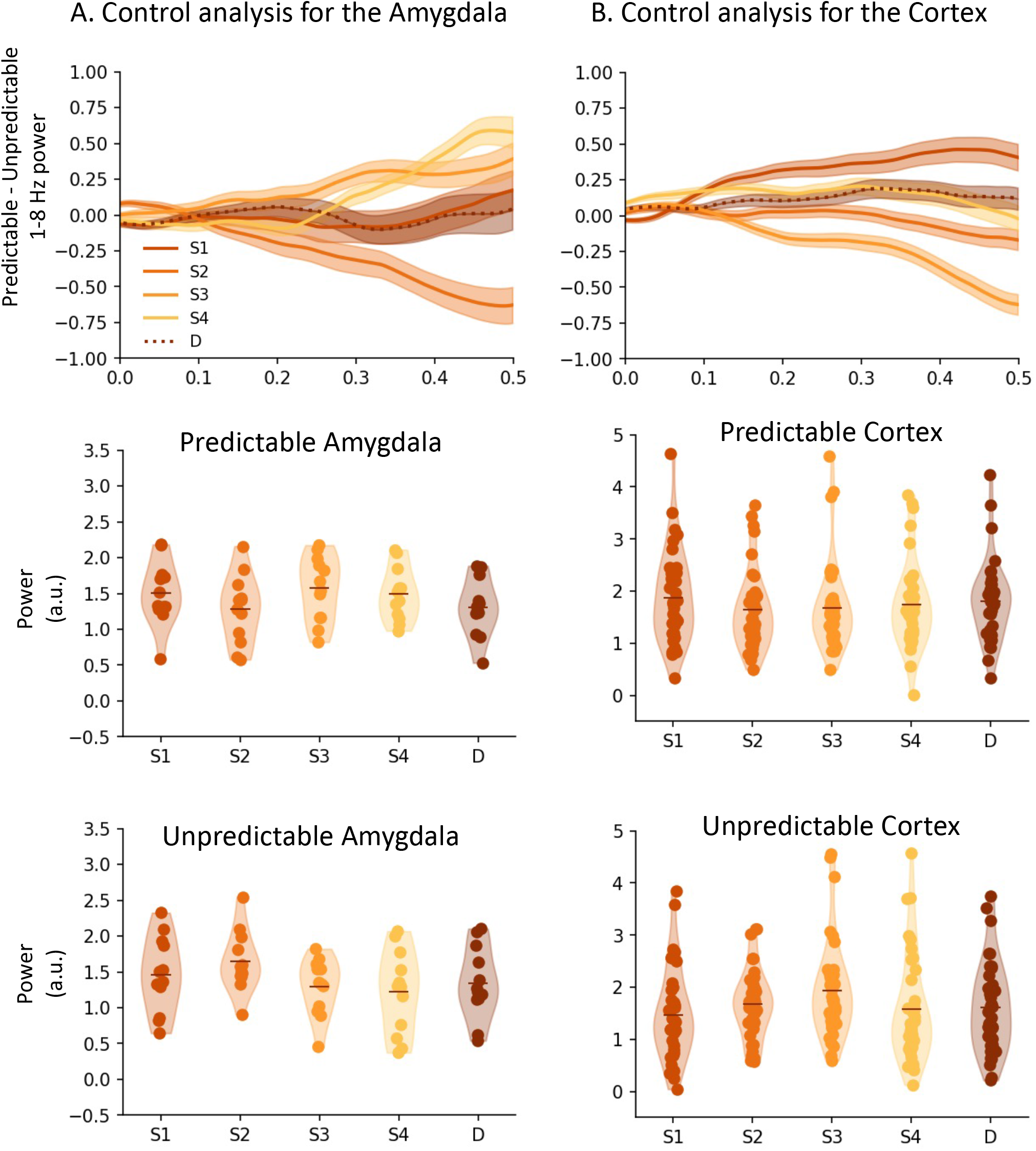
Control analysis for the evolution of low frequency power in the amygdala and cortex, as the sequence unfolds. Neither the 1-8 Hz power over time, nor a data-driven low power definition showed any significant effect of sequence.

